# Reverse genetic screen reveals that Il34 facilitates yolk sac macrophage distribution and seeding of the brain

**DOI:** 10.1101/406553

**Authors:** Laura E. Kuil, Nynke Oosterhof, Samuël N. Geurts, Herma C. van der Linde, Erik Meijering, Tjakko J. van Ham

## Abstract

Microglia are brain resident macrophages, which have specialized functions important in brain development and in disease. They colonize the brain in early embryonic stages, but few factors that drive the migration of yolk sac macrophages into the embryonic brain, or regulate their acquisition of specialized properties are currently known.

Here, we present a CRISPR/Cas9-based in vivo reverse genetic screening pipeline to identify new regulators important for microglia development using zebrafish. Zebrafish larvae are particularly suitable due to their external development, transparency, high fecundity and conserved microglia features. We targeted putative microglia regulators, including signature genes and non-cell autonomous factors, by Cas9/gRNA-complex injections, followed by neutral red-based visualization of microglia. Microglia were quantified automatically in 3-day-old larvae using a software tool we called SpotNGlia. We identified that loss of function of the zebrafish homolog of the colony stimulating factor 1 receptor (CSF1R) ligand IL34, caused strongly reduced microglia numbers in early development. Previous studies on the role of the IL34 on microglia development in vivo were ambiguous. Our data, and a concurrent paper, show that in zebrafish, *il34* is required during the earliest seeding of the brain by microglia progenitors. Our data also indicate that Il34 is required for distribution to other organs.

Previously, we showed that *csf1ra* and *csf1rb* double mutant zebrafish have no microglia. As there is a moderate effect of *il34* on microglia development, relative to the effect *csf1r*, additional Csf1r-dependent signalling may be needed for establishment of the microglia network. In all, we identified *il34* as a modifier of microglia colonization, by affecting distribution of yolk sac macrophages to target organs, validating our reverse genetic screening pipeline in zebrafish which can be used for the identification of additional regulators of microglia development.

## INTRODUCTION

Tissue macrophages, in addition to their immunological roles, modulate organogenesis and exhibit organ-specific regulatory properties that are thought to affect virtually all organs in vertebrates (1, 2). Microglia are the brain’s resident macrophages, which have roles in brain development and homeostasis. Described functions of microglia include the removal of dead cells and debris, modulation of neuronal connectivity by synaptic pruning and maintenance of myelin-producing cells (3-6). Defects in microglia function have been implicated in neurodevelopmental disorders such as autism spectrum disorder (ASD)(3). Pathogenic variants in genes thought to primarily affect microglia cause rare white matter disorders including Nasu-Hakola disease and adult onset leukoencephalopathy with axonal spheroids (ALSP), which may be caused by loss of microglia activity (7-10). In line with this, there is accumulating evidence that replenishing brain myeloid cells by hematopoietic cell transplantation (HCT) has powerful therapeutic potential in leukodystrophy and metabolic diseases affecting the brain, and better understanding the molecular regulation of brain colonization by microglia could lead to ways to facilitate this (11-13). However, the exact genes and mechanisms underlying the emergence of microglia in the brain and acquisition of their functional properties are still poorly understood.

Microglia originate from macrophage progenitors in the embryonic yolk sac that colonize the brain during early embryonic development (14, 15). Once they arrive in the brain, they acquire a highly ramified morphology, proliferate extensively and form a brain-wide network with non-overlapping territories (16). The transition from yolk sac macrophage to mature microglia or other tissue resident macrophages involves several differentiation stages characterized by distinct transcriptional profiles (17, 18). The progression through these transcriptional states is synchronised with, and most likely driven by, the different stages of brain development as microglia gene expression is highly sensitive to changes in the microenvironment and tissue macrophage identity is mostly determined by the host environment (17, 19-21). For the majority of the genes specifically expressed in microglia the function is still unknown, and as many of these genes are rapidly downregulated when they are taken out of the brain, it is difficult to study their functions in vitro (22, 23). In mammals, microglia development is relatively inaccessible to study, as progenitors emerge during development in utero. Despite progress in identifying methods to recreate microglia-like cells in vitro, improved understanding of their ontogeny is needed to guide in vitro efforts (24, 25). Therefore, identification of the functions of genes affecting microglia development could provide valuable insights into regulation of microglia development and function in vivo.

Zebrafish embryos are relatively small, transparent, are relatively easy to manipulate genetically and develop ex-utero, which makes them highly suitable for in vivo genetic studies (26). We recently showed that microglia gene expression is well conserved between zebrafish and mammals and that, as shown in mice, loss of the two zebrafish homologs of the colony-stimulating factor 1 receptor (Csf1ra and Csf1rb) leads to absence of microglia (10, 27-29). Phenotype-driven, forward genetic screens in zebrafish have identified several microglia mutants with a defect in microglia development or function. Processes affected in these mutants include hematopoiesis, regulation of inflammation, phosphate transport and lysosomal regulation, which implies that these various processes are all critical for microglia development and function (30-34). However, such forward genetic screens are laborious and relatively low-throughput. A candidate-driven reverse genetic screening approach could lead to identification of additional genes important for microglia. The CRISPR/Cas9-system can be used to create insertions or deletions (indels) in target genes via the repair of Cas9-induced double strand breaks by error-prone non-homologous end joining (NHEJ) (35). Injection of gene specific guide RNAs (gRNAs) and Cas9 mRNA, can lead to gene disruption sufficiently effective to allow small-scale reverse genetic screening, for example to identify new genes involved in electrical synapse formation (36). Alternatively, active Cas9-gRNA ribonucleoprotein complexes injected into fertilized zebrafish oocytes can more efficiently induce indels in target genes and the resulting genetic mosaic zebrafish can phenocopy existing loss-of-function mutants (CRISPants) (37, 38).

Here, we present a scalable CRISPR/Cas9-based reverse genetic screening pipeline in zebrafish to identify important genetic microglia regulators using zebrafish. In zebrafish larvae, microglia can be visualized by the vital dye neutral red, which is actively taken up by phagocytosis and has been used as an effective readout in forward genetic screens (15, 30-32). We developed an image quantification tool, SpotNGlia, to automatically detect the brain boundaries and count neutral red-positive (NR+) microglia. Out of the 20 putative microglia regulators we targeted by CRISPR/Cas9-mediated reverse genetics, disruption of *interleukin 34* (*il34*) showed the strongest reduction in microglia numbers in developing zebrafish larvae. In mammals, Il34 is one of two ligands of the microglia regulator CSF1R. Further analysis in stable *il34* mutants revealed that *il34* is mainly important for the recruitment of microglia to the brain, and likely other tissue resident macrophage populations, including Langerhans cells, to their target organs. Thus, we here present a scalable reverse genetic screening pipeline to identify additional new regulators important for microglia development and function.

## RESULTS

### CRISPants phenocopy existing mutants with microglia developmental defects

Loss of one of several key macrophage regulators, including *Spi1* (encoding PU.1), *Irf8* and *Csf1r*, and their zebrafish homologs *spi1b* (Pu.1), *csf1ra* and *csf1rb*, and *irf8*, leads to defects in microglia development (15, 28, 39-43). To investigate whether Cas9-gRNA ribonucleoprotein complexes (RNPs) targeting these regulators can be used to induce mutant microglia phenotypes directly, we injected zebrafish oocytes with RNPs targeting either *csf1ra* or *spi1b.* To assess whether CRISPR/Cas9-based targeting of those genes affects microglia development we determined microglia numbers by neutral red (NR) staining at 3 days post fertilization (dpf). At this time point, microglia have just colonized the optic tectum, are highly phagocytic and have low proliferative activity, which makes it an ideal time point to identify genes required for the earliest steps of microglia development (15, 44). We quantified NR+ microglia in *csf1ra* CRISPants, in controls and in *csf1ra* loss-of-function mutants found in an ENU mutagenic screen (hereafter called *csf1ra*^-/-^)(45). Similar to *csf1ra*^*-/-*^ mutants, *csf1ra* CRISPants showed an 80% reduction in the number of NR+ microglia compared to controls suggesting highly effective targeting in F0 injected embryos (Fig 1A). To assess the targeting efficiency of the *csf1ra* gene we performed Sanger sequencing of the targeted locus of a small pool of *csf1ra* CRISPants and calculated the spectrum and frequency of indels in the *csf1ra* gene using TIDE (tracking indels by decomposition) software (46). The mutagenic efficiency was >90%, showing efficient mutagenesis (Fig 1B). Similarly, *spi1b* CRISPants showed a strong reduction in the number of microglia and 65-95% mutagenic efficiency (Fig 1C, D). This shows that CRISPR/Cas9-based mutagenesis can be used to reproduce mutant microglia phenotypes in Cas9-gRNA RNP injected zebrafish larvae.

**Fig 1.**
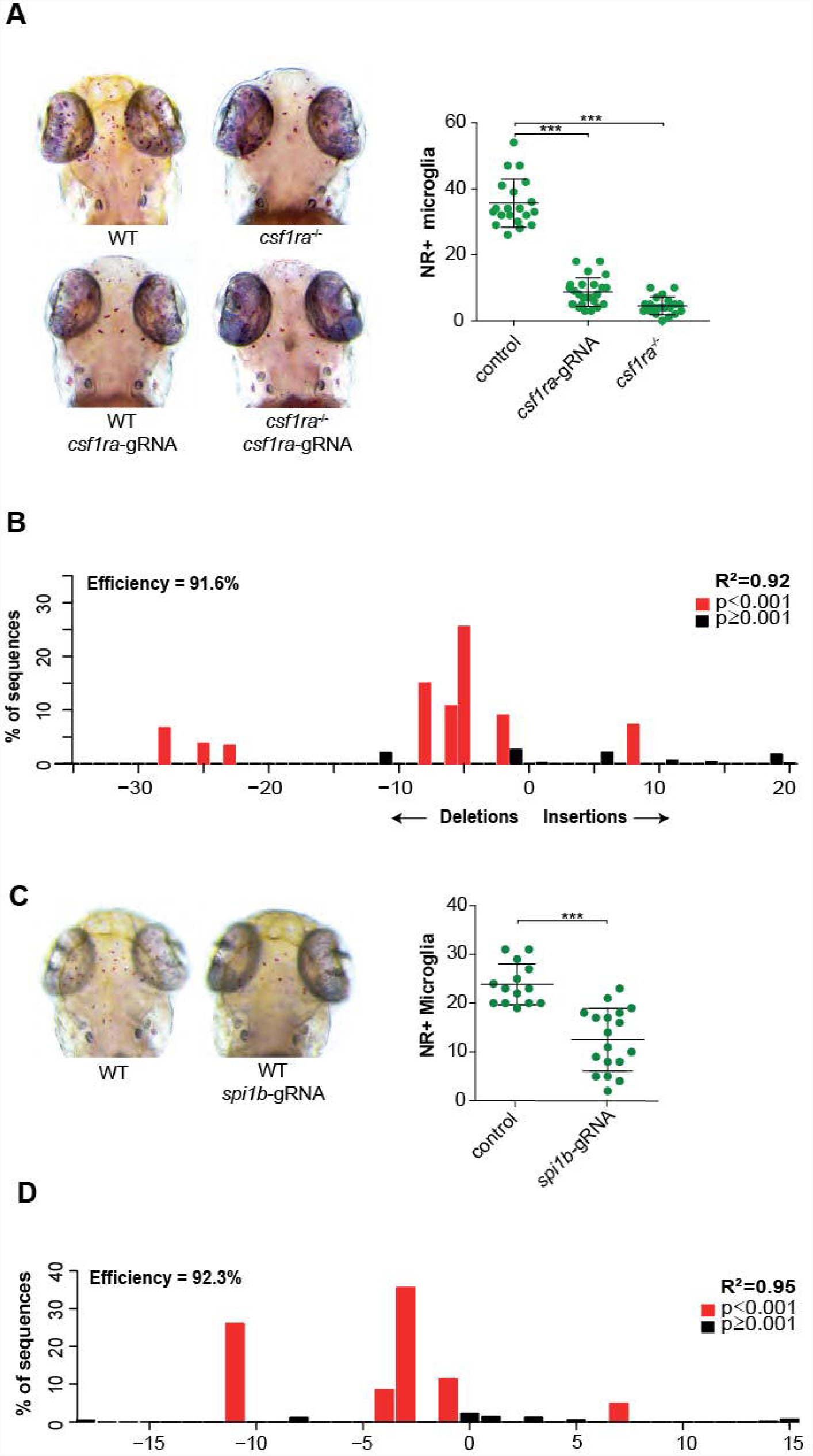
*csf1r* CRISPants phenocopy existing *csf1r* microglia mutants. (A) Neutral red (NR+) images and quantification of WT, *csf1ra*^-/-^ and *csf1ra* CRISPant zebrafish larvae at 3 dpf. (B) Indel spectrum of a pool of *csf1ra* CRISPants calculated by tide. (C) Neutral red images and quantification of WT, and *spi1b* CRISPant zebrafish larvae at 3 dpf. (D) Indel spectrum of a representative individual *spi1b* CRISPant calculated by tide. R^2^ value represents reliability of the de indelspectrum. *** p < 0.001.

### SpotNGlia semi-automatically counts microglia numbers

Manual quantification of NR+ microglia, across z-stack images, is time-consuming and can be subjective. To standardize and speed up quantification, we developed a software tool, SpotNGlia, that automatically counts NR+ microglia in the optic tectum where most microglia are located at 3 dpf. The SpotNGlia tool aligns stacked images of stained zebrafish larvae taken at different axial positions and blends the images into a single 2D image in which all NR+ cells are in focus (Fig 2A). Next, the images are segmented by using polar transformation and dynamic programming to identify the edges of the optic tectum. Finally, NR+ cells are detected and counted by a spot detection technique based on multiscale wavelet products (47). To test the SpotNGlia software tool, we created and manually annotated a dataset with representative z-stack images of 50 neutral red stained zebrafish larvae. To assess the accuracy of brain segmentation, Jaccard and Dice indices were determined, revealing indices of 0.86 (Jaccard) and 0.93 (Dice)(Fig 2B, C). To assess the accuracy of microglia detection we determined the precision, recall and F1 scores of the computed annotation, resulting in average scores of 0.85, 0.91 and 0.87, respectively (Fig 2B,C,D). These results indicate that SpotNGlia is able to automatically identify the boundaries of the midbrain region, and the microglia within that region, in the vast majority of cases. To correct manually for those instances where brain segmentation and microglia detection were not completely accurate -as determined by visual inspection, our tool offers the possibility of post-hoc correction. In our experiments we have found that SpotNGlia results in about 80% reduction in the time it takes to quantify NR+ microglia numbers. In all, this indicates that SpotNGlia is a powerful tool for fast quantification of NR+ microglia numbers to assist in identifying novel genes important for generation of functional microglia.

**Fig 2.**
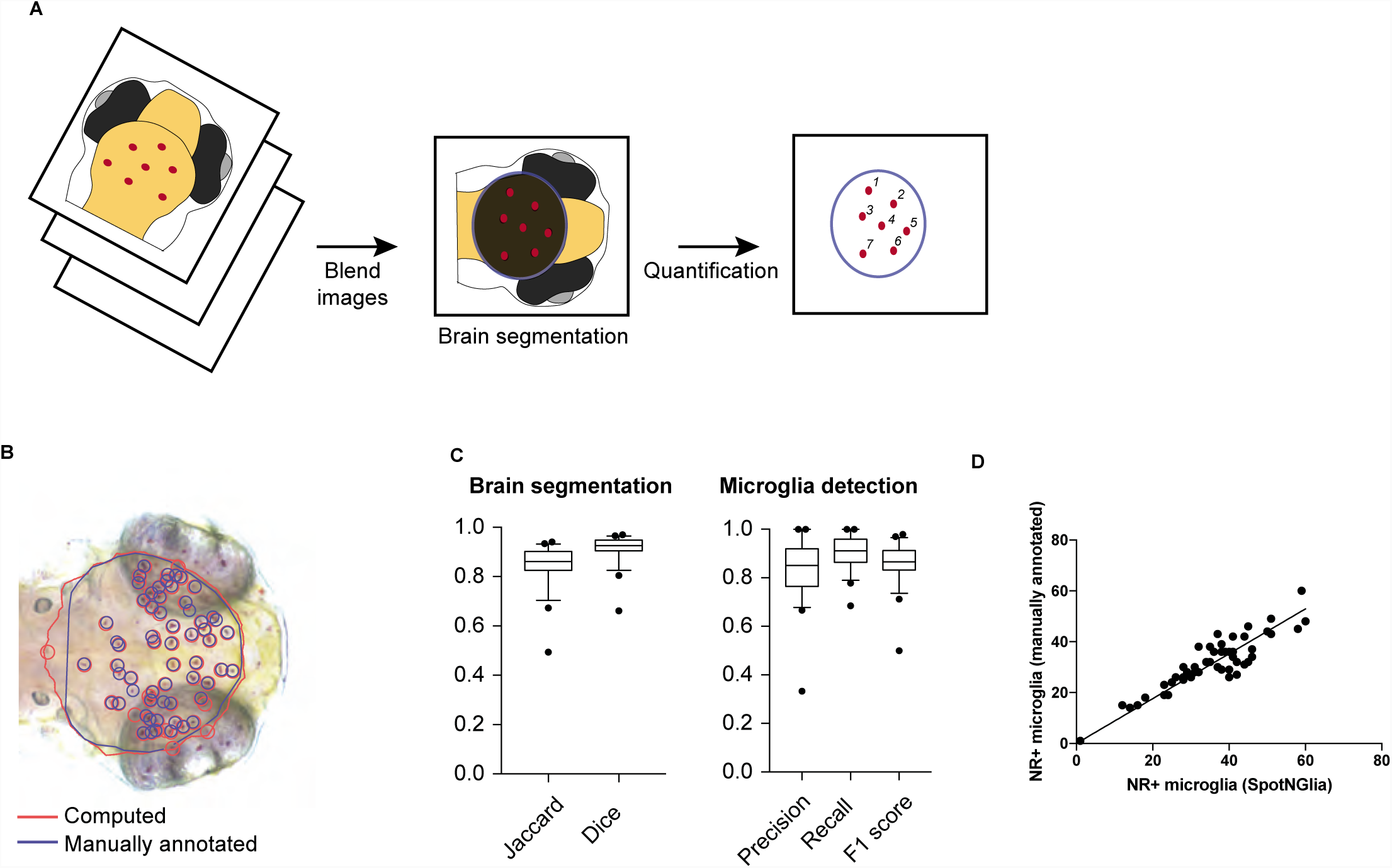
SpotNGlia semi-automatically counts microglia numbers. (A) Schematic representation of SpotNGlia analysis pipeline. (B) SpotNGlia output of test dataset with both manual (blue) and automated (red) brain segmention and NR+ microglia annotation. (C) Boxplots showing Jaccard and Dice indices for accuracy of brain segmentation and F1, precision and recall scores for the accuracy of NR+ microglia annotation. (D) Correlation between manually and automated microglia quantification after manual correction for segmented brain area.

### Reverse genetic screen reveals zebrafish Il34 as a regulator of microglia development

To identify new microglia regulators using direct CRISPR/Cas9-targeting and microglia phenotyping by SpotNGlia, we targeted 20 candidate genes individually. These genes were selected based on either our recently identified zebrafish microglia transcriptome (e.g. *slco2b1, hcst*/*dap10* and *mrc1b*), microglia expressed genes with a connection to brain disease (*e.g. usp18*), or genes which could affect microglia in a non-cell autonomous manner (CSF1R ligand encoding genes *il34, csf1a* and *csf1b*) (Fig 3A, Table S1)(27). Next, gRNAs were designed to effectively target these genes in one of their first exons. Cas9-gRNA RNPs targeting candidate genes were injected in fertilized oocytes, after which they were NR stained at 3 dpf, phenotyped and genotyped by Sanger sequencing followed by indel decomposition using TIDE (Table S1)(46). We did not observe obvious signs of developmental delay, morphological abnormalities or increased mortality upon Cas9-gRNA RNP injections, indicating that the observed microglia phenotypes were not due to Cas9-gRNA toxicity. The gRNAs for 6 of the targeted genes caused a significant reduction in the number of NR+ microglia (Fig 3A). The largest decrease in NR+ numbers was observed in embryos in which the zebrafish homolog of interleukin 34 (IL34) was targeted (Fig 3A, B)(48).

**Fig 3.**
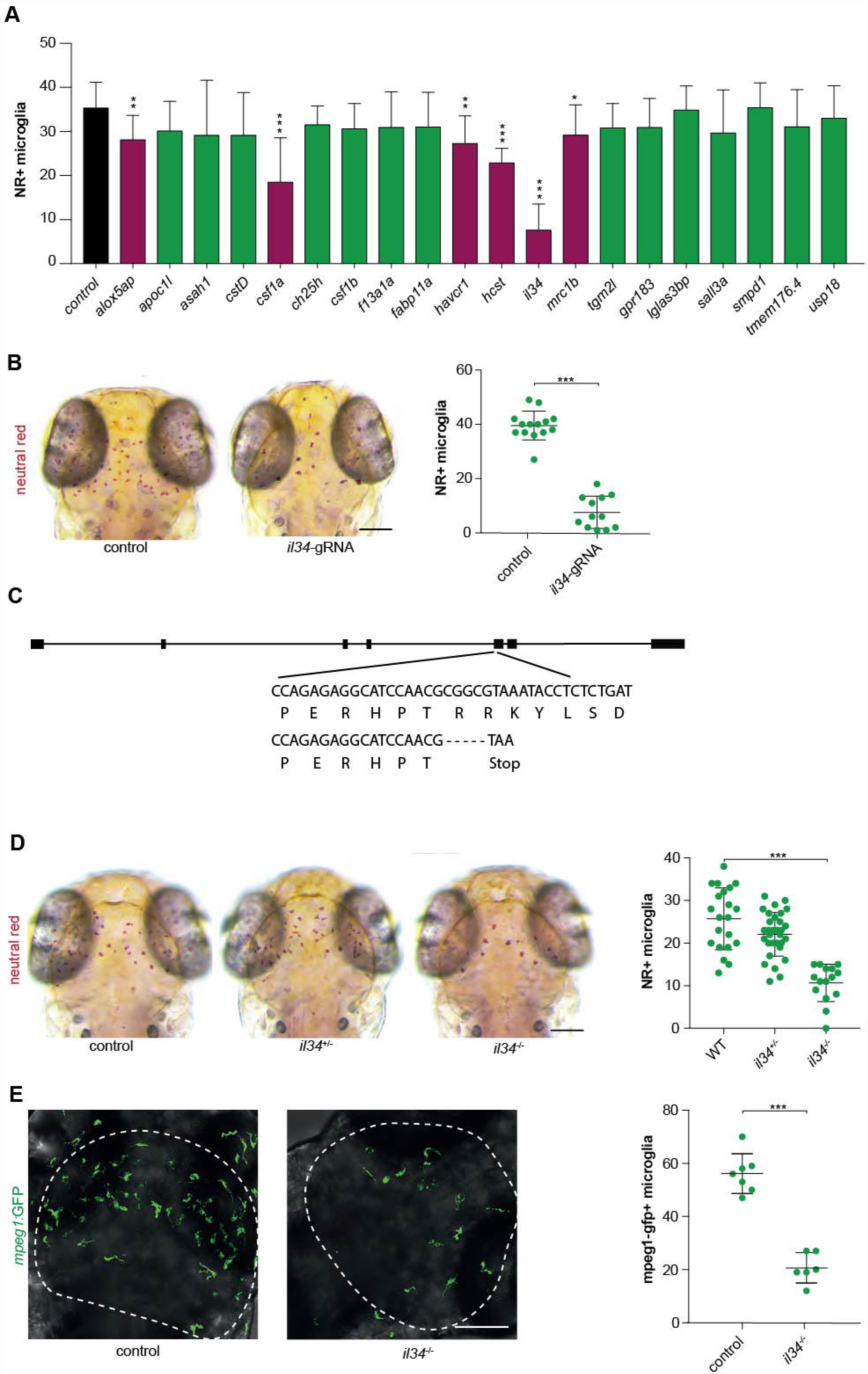
Reverse genetic screen reveals zebrafish *il34* as a regulator of microglia development. (A) Accumulated data from all gRNA injections showing the number of NR+ microglia as quantified with SpotNGlia. Red bars represent genes which showed a significant reduction in microglia numbers upon CRISPR/Cas9-based targeting. (B) NR+ microglia numbers in 3 dpf zebrafish larvae injected with gRNA-Cas9 RNPs targeting *il34*. (C) −5 bp deletion in exon 1 of *il34* directly introduces a stop codon (D) NR+ microglia numbers in *il34* mutants with a premature stop codon in exon 5 and their littermates at 3 dpf. (E) GFP+ microglia in the optic tecti (dotted line) of 3 dpf *il34* mutants and controls and quantification. * p < 0.05, ** p < 0.01,*** p < 0.001. Scale bar represents 100 µm.

To validate our approach and confirm that this microglia phenotype is caused by loss of *il34* function, we generated a premature stop codon in exon 5 of the *il34* gene. Neutral red labelling of homozygous *il34* mutants at 3 dpf revealed a ∽60% reduction in NR+ microglia compared to wildtype siblings, suggesting this is a loss of function allele (Fig 3C). Similarly, live imaging of GFP expressing microglia (GFP+), driven by the *mpeg1* promoter, in the optic tecti of *il34* mutants showed lowered microglia numbers compared to controls (Fig 3D). In mice, *Il34* knockout led to slightly different outcomes, causing, in one study, lowered microglia numbers already in early postnatal development that remained low into adulthood and, in another study, only reduced adult microglia numbers (49, 50). Therefore, the precise role of Il34 in early microglia development remains ambiguous. In addition, the precise role of Il34 in adult microglia has not been described yet (49, 50). Our results are consistent with an evolutionary conserved role for *Il34* in early microglia development (49). This is further supported by a concurrent study where, using another premature stop mutation in *il34*, the authors showed that mutation of *il34* leads to a similar reduction in microglia numbers at the same developmental stage (Wu et al., 2018, Dev Cell).

### Il34 facilitates the distribution of macrophages, without affecting their proliferation

In mice, tissue resident macrophages of the skin, known as Langerhans cells (LCs), are highly dependent on IL34/CSF1R-signaling for their maintenance and self-renewal (49-51). We therefore hypothesized that Il34 in zebrafish might regulate the proliferative expansion of microglia, similar to LCs in mice, leading to the lower microglia numbers we observed. Microglia numbers increase sharply after 3 dpf, and to determine whether microglia numbers remained lower over time we quantified NR+ microglia also at 5 dpf (Fig 4A). Surprisingly, compared to 3 dpf, microglia numbers in *il34*-/- mutants were closer to those of controls at 5 dpf (∽30% reduction 5 dpf vs ∽60% reduction at 3dpf). To determine whether the increase in numbers was due to the continuation of seeding the brain or proliferative expansion we performed EdU pulse labelling between 3 and 4 dpf. EdU/L-plastin double labelling showed reduced microglia and reduced Edu+ microglia, but the fraction of EdU+ microglia did not differ between *il34* mutants and controls (Fig 4B). Thus, loss of *il34* does not change the proliferative fraction of microglia, therefore the decreased microglia numbers are unlikely explained by a defect in proliferation. Since the decrease in microglia numbers in *il34* mutants compared to controls was largest at 3 dpf, Il34 likely affects microglia progenitors preceding brain colonization. Indeed, Wu and colleagues show that Il34 deficiency causes impaired colonization by failing to attract progenitors to enter the brain in a Csf1ra-dependent mechanism (Wu et al., 2018). We used live-imaging to visualize *mpeg1*-GFP+ yolk sac macrophages (YSMs), the progenitors of microglia and many other macrophages. At 2 dpf, YSM numbers and morphology were not different between *il34* mutants and controls (Fig 4C). Thus, reduced microglia numbers are likely not attributed to reduced progenitor pool size. Therefore impaired migration of *il34* deficient macrophages towards the brain could explain the lower microglia numbers. Imaging in the rostral/head region at 2 dpf showed an >80% decrease in the number of macrophages, suggesting that *il34* is indeed involved in the recruitment of YSMs to the brain (Fig 4D). To determine whether this effect is exclusive to microglia, we determined the fraction of total macrophages that was found in the head or in the trunk region at 3 dpf. This showed again an ∽80% reduced infiltration of microglia progenitors in the brain in *il34* mutants compared to controls. Colonization of the trunk was also decreased in *il34* mutants compared to controls, but to a lesser extend (25% reduction)(Fig 4E-F). Therefore, *il34* appears particularly important for the recruitment of YSMs towards the brain. In addition, it also affects the distribution into the trunk region, possibly analogous to the effect of IL34 on the maintenance and development of LCs, as described in mice (49-51).

**Fig 4.**
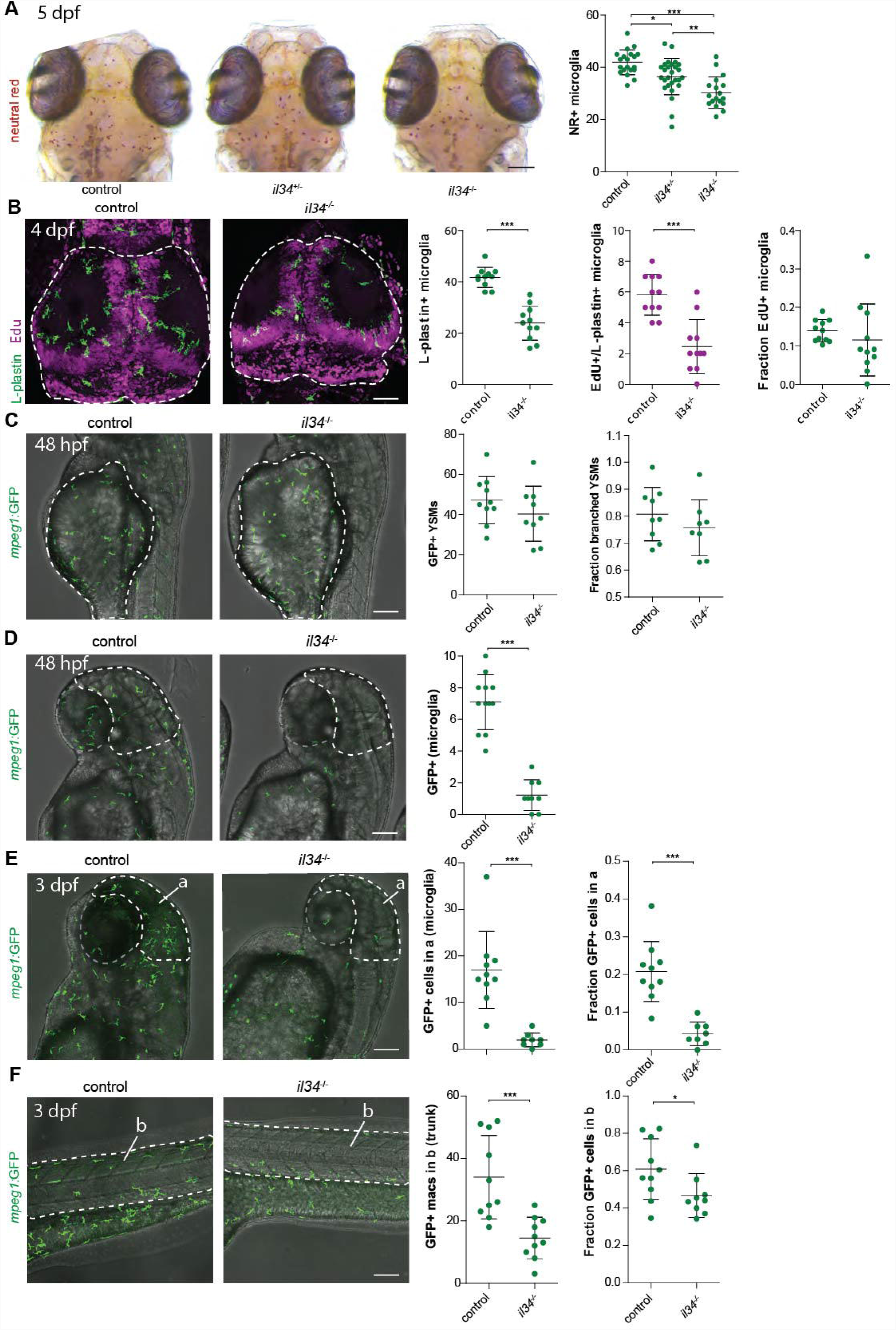
Il34 does not affect proliferation but the distribution of YSMs to target organs. (A) NR+ microglia numbers in *il34* mutants and their siblings at 5 dpf. (B) EdU/Lplastin staining of microglia in the optic tecti (dotted line) of 4 dpf *il34* mutants and controls and quantification of microglia numbers, EdU+ microglia numbers and the fraction of EdU+ microglia among total numbers. (C) In vivo imaging of GFP+ macrophages located on the yolk sac (dotted line) in *il34* mutants and controls, transgenic for *mpeg1-GFP*, and, quantification at 48 hpf. YSMs with more than 1 protrusion were counted as branched YSMs. (D) In vivo imaging of mpeg1:GFP+ macrophages located in the head region (dotted line) in *il34* mutants and controls and its quantification at 48 hpf. (E) In vivo imaging of GFP+ macrophages located in the head region (dotted line) in *il34* mutants and controls and its quantification at 3 dpf. (F) In vivo imaging of mpeg1:GFP+ macrophages located in the tail (dotted line) in *il34* mutants and controls and its quantification. Scale bar represents 100 µm. * p < 0.05, ** p < 0.01, *** p < 0.001

## DISCUSSION

In this study, we developed a scalable CRISPR/Cas9-based reverse genetic screening pipeline using semi-automated image quantification to identify new regulators of microglia biology using zebrafish embryos. We showed that direct genetic targeting of known microglia regulators including *csf1ra* and *spi1b* by Cas9/gRNA injections in zebrafish embryos phenocopies previously identified microglia mutants. We next developed a software tool (SpotNGlia) that allows for automated phenotyping by quantification of neutral red positive microglia. As zebrafish are well suited for in vivo drug discovery, our strategy could potentially also be used to identify small molecules affecting microglia development (52). Using this pipeline, we here tested 20 candidate genes for a role in microglia development and found 6 genes significantly affecting microglia numbers when mutated. Loss of *il34* function caused the largest decrease in microglia numbers, which we confirmed by analysis of stable mutants. Furthermore, we uncovered *il34* as a regulator of distribution of tissue macrophages, needed to recruit microglia progenitors to the brain.

Even though we here examined 20 genes, there are several ways to increase the throughput of our screening strategy. First, mounting of the injected zebrafish larvae and subsequent image acquisition are the most time-consuming parts of our pipeline. Neutral red stained larvae were manually embedded in low melting point agarose before imaging, which restricts the number of animals that can be screened per day. Automated imaging systems that can load zebrafish larvae from liquid medium in multi-well plates and image them in the orientation of interest in glass capillaries could overcome this hurdle (53). Together with the SpotNGlia tool this would permit a significantly increased screening throughput and efficiency. Additionally, we aimed to achieve maximal CRISPR/Cas9 mutagenic efficiency for individual genes of interest, and therefore targeted individual genes. Shah et al. previously reported a strategy where pools of up to 8 gRNAs are injected simultaneously to target multiple genes at once (54), which could lead to reduced targeting efficiency of the individual gRNAs. Although a pooling strategy could significantly increase the number of genes that can be screened, we observed that, especially for genes with a relatively subtle microglia phenotype, a high mutagenic efficiency increases the chance of detecting the phenotype. Additionally, due to the clonal nature of hematopoietic progenitors, including yolk sac macrophages, a high targeting efficiency is likely required, because non-targeted cells could expand and compensate for mutated cells.

IL34 is one of two ligands of the colony stimulating factor 1 receptor (CSF1R), a main regulator of development of the macrophage lineage (55). Even though adult *Il34* deficient mice have fewer microglia, and no Langerhans cells, the precise role of IL34 in microglia development is unclear. Wang and colleagues, showed that neonatal *Il34*-/- mice have lower microglia numbers, whereas Greter et al showed normal microglia numbers in *Il34*-/- mice throughout embryonic development (49, 50). The exact function of Il34 in microglia development in vivo, and how this may differ from Csf1, remains therefore ambiguous. These discrepancies could be attributed to factors such as genetic background, or slightly different methods leading to different interpretations regarding the role of Il34 in embryonic and early postnatal microglia numbers (49, 50).

Our data revealed a ∽60% reduction in microglia numbers in *il34* mutant larvae at 3 dpf, indicating that *il34* is required for early microglia development in zebrafish. We show that upon arrival in the brain, between 3 and 5 dpf, microglia numbers increase by proliferation in both controls and *il34* mutants, suggesting that the proliferative capacity of microglia is not affected by the loss of *il34*. In addition, microglia progenitor numbers on the yolk sac were not affected by *il34* deficiency, indicating that there is a defect in the colonization of the embryonic brain by macrophages, likely due to a failure to attract progenitors expressing Csf1ra and/or Csf1rb. Consistent with this, analysis of migration towards the brain at both 2 and 3 dpf showed much fewer microglia colonized the brains of *il34* deficient larvae. Our findings are consistent with a concurrent manuscript, where the authors show that nervous system expression of Il34 can attract microglia progenitors to migrate into the brain by the Il34/Csf1 receptor Csf1ra (Wu et al., 2018). However, we additionally found that distribution of *il34* mutant macrophages into trunk regions was reduced, indicating that the effect of Il34 is not limited to microglia, but also affects the migration and colonization of other macrophages.

As we previously showed that mutants for both receptors, Csf1ra and Csf1rb, lack all microglia, in contrast to Csf1ra mutants, which have fewer microglia only in early development, the expansion of microglia progenitors following colonization of the brain is likely regulated by other, possibly compensatory or redundant, factors, including through CSF1 homologs *csf1a* and *csf1b* (10, 29). Indeed, in *il34* mutants, many precursors continue to arrive in the brain at 3 dpf, which start to proliferate and reach 70% of control levels at 5 dpf. As well, numerous studies have shown that CSF1 activation of CSF1R drives macrophage proliferation (56), and *Csf1* and *Il34* were both found to be expressed in the brain, although in non-overlapping regions (49, 57). Thus, we find that *il34* is a critical, non-cell autonomously regulator of seeding of the brain by microglia progenitors, but likely not their proliferation, which also facilitates the distribution of macrophages to other organs.

In conclusion, we here present a scalable reverse genetic screening method for the identification of novel regulators of microglia development and function. Microglia are key players in brain disease and there is strong evidence that microglia defects can be a primary cause of brain disease (7-10). Replenishing microglia, for example by hematopoietic stem cell transplantation (HCST) can provide therapeutic benefit in human brain diseases. Better understanding of microglia development and acquisition of their specific cell fate in vivo, could lead to improved strategies to replace defective microglia. However, the mechanisms and genes regulating microglia development and function are still largely unknown. Therefore, better understanding of microglial gene functions could be a valuable step in the elucidation of mechanisms underlying microglial biology. As zebrafish larvae have proven their suitability for drug discovery, SpotNGlia automated analysis software in combination with automated imaging systems could also be used to screen for compounds affecting microglia (58). In all, we identified *il34* as a regulator of tissue resident macrophage distribution, primarily affecting macrophage colonization of the zebrafish embryonic brain by affecting the recruitment of YSMs to target organs including the brain. Our reverse genetic screening pipeline can be used to address genetic regulation of microglia development and function, and identify regulators essential to generate functional microglia in vivo.

## MATERIALS AND METHODS

### Fish care

For all experiments Tg(*mpeg1*:*EGFP*) fish expressing GFP under the control of the *mpeg1* promotor or Tg(*Neuro*-*GAL4, UAS*:*nsfB*-*mCherry, mpeg1*:*EGFP*) with neuronal specific nitroreductase expression, transgenic zebrafish lines were used (59). Zebrafish embryos were kept at 28°C on a 14h/10h light-dark cycle in HEPES-buffered E3 medium. At 24 hpf 0.003% 1-phenyl 2-thiourea (PTU) was added to prevent pigmentation.

### sgRNA synthesis

To design sgRNAs the online program CRISPRscan (www.crisprscan.org) was used (Moreno-Mateos et al., 2015). The gRNAs were designed to target exons, except for exon 1, to be as close as possible to the transcription start site and to have no predicted off-target effects. The sgRNAs were generated from primer dimers, containing a minimal T7 RNA polymerase promoter. To generate primer-dimers the FastStart™ High Fidelity PCR System from Sigma was used. A solution was prepared containing 1 mM forward sgRNA oligo, 1 mM reverse oligo consisting of 20 nt overlap with sgRNA oligo and the Cas9-binding part, 0.8 mM dNTPS, 1x FastStart Buffer and 6.25 U / µL FastStart Taq polymerase in 20 µL total volume. Annealed DNA oligo dimers were generated by denaturation at 95°C for 5 minutes followed by annealing by reducing the temperature by 1°C per second during 20 seconds to 75°C f and extension at 72°C for 10 minutes. The gRNAs were synthetized from annealed DNA oligo’s, containing a minimal T7 RNA polymerase promoter, with the mMESSAGE mMACHINE T7 ULTRA Transcription Kit (Invitrogen) according to the manufacturer’s instructions.

### Cas9/gRNA complex injections into zebrafish larvae

The SP-Cas9 plasmid used for the production of Cas9 protein was a gift from Niels Geijsen (Addgene plasmid #62731) (60). Cas9 nuclease was synthetized as described (60). 600-900 ng of gRNA was mixed with 4 ng of Cas9 protein to form active gRNA-Cas9 RNPs. Next, 0.4 µL of 0.5% Phenol red (Sigma) and the volume was adjusted with 300 mM KCl to a total volume of 6 µL. Approximately 1 nL of the mix was injected in fertilized zebrafish oocytes. For the creation of the *il34* mutant line CRISPants were grown to adulthood and outcrossed to the AB background, and Sanger sequencing was used to identify mutations.

### Neutral red staining and imaging

To label microglia, 3 dpf larvae were incubated in E3 medium containing neutral red (Sigma) (2.5 ug/ml) to for 2 hours at 28 °C, after which they were rinsed with E3 medium containing 0.003% PTU. Larvae were anaesthetized with 0.016% MS-222 and embedded in 1.8% low melting point agarose in E3 with the dorsal side facing upwards. Serial images (3-6) in the z-plane were acquired with a Leica M165 FC microscope using the 12x dry objective and a Leica DFC550 camera.

### Larvae genotyping (Sanger sequencing)

#### Lysis

Zebrafish larvae were euthanized and placed in single tubes containing 100 µL lysis Buffer (0.3% 1M KCl, 1% 1M TrisHCl pH 9.0, 0.1% Triton, 0.15 mg/mL Proteinase K) per larva. The mix was incubated at 55 °C for 10 minutes and 95 °C for 10 minutes. The lysate was centrifuged for 5 to 10 minutes at 4000 rpm, and 1 µl was used for PCR.

#### Sanger sequencing to determine CRISPR/Cas9 targeting efficiency

For Sanger sequencing 500 bp long PCR products were obtained. For the sequencing reaction BigDye® Terminator v3.1 Cycle Sequencing Kit from Applied Biosystems was used. The product was placed on Sephadex® columns (Sigma) and centrifuged at 910 rcf for 5 minutes. The ABI 3130 genetic analyzer from Applied Biosystems was used for sanger sequencing. To assess the indel spectrum and frequencies at the target locus we used the program TIDE developed by the Netherlands Cancer Institute (NKI)(46).

### SpotNGlia

#### Preprocessing

Images acquired from neutral red labelled larvae (n=50) were used to optimize the algorithm. For each larva, 3-6 images were taken at different depths of focus. Color channels were realigned by finding the translation that maximizes the correlation coefficient (61). To remove the background the triangle thresholding method was used (62). Next, we generated an all-in-focus image with extended depth of field (63).

#### Brain segmentation

The orientation of the fish was determined by maximizing the correlation coefficient between the image and a mirrored version of itself, yielding the larvae’s rotation angle. The translation parameters were found by directly correlating the image to a template image, which was established by averaging multiple aligned fish. Because of its near-circular shape, the optic tectum was segmented by performing a polar transformation after which the edges of the optic tectum were found by using Dijkstra’s algorithm (64, 65). The brain edge becomes an approximately straight line in polar coordinates if it is transformed with respect to the center of the optic tectum which we obtained from the template image. To make it applicable for the shortest path algorithm, the image was correlated with a small image, similar to the average appearance of the brain edge in the polar image. Also a priori information of the training set was used to exclude locations where the brain edge cannot be. After Dijkstra’s algorithm was applied the found path was transformed back resulting in the brain edge coordinates.

#### Microglia detection

To identify neutral red-positive (NR+) microglia a multiscale wavelets product was computed on the green channel of the image, which contains the highest contrast for the NR signal (47). Multiple smoothed images from a single fish image were produced with increasing spatial scale. Subtracting adjacent smoothed images resulted in subband images containing different scales of detail present in the image. A product of subband images in the range of the microglia spot size was performed to obtain an image with only high values at the location of the spots, i.e. the multiscale wavelet product. A threshold on the multiproduct image was applied to obtain a binary image to determine the spots. The identified spots were discriminated further on typical color and size obtained from the training set, resulting in accurate quantification of microglia numbers.

### Immunofluorescence staining

Immunohistochemistry was performed as described (66, 67). Briefly, larvae were fixed in 4% PFA at 4°C overnight. Subsequently, dehydrated to 100% MeOH and stored at −20°C for at least 12 hours, and rehydrated to PBS. Followed by incubation for three hours in blocking buffer (10% goat serum, 1% Triton X-100 (Tx100), 1% BSA, 0.1% Tween-20 in PBS) at 4°C was followed by incubation in primary antibody buffer at 4°C overnight. Larvae were washed in 10% goat serum 1% Tx100 in PBS and PBS containing 1% TX100 for a few hours, followed by incubation in secondary antibody buffer at 4°C overnight. Primary antibody buffer: 1% goat serum, 0.8% Tx100, 1% BSA, 0.1% Tween-20 in PBS. Secondary antibody buffer: 0.8% goat serum, 1% BSA and PBS containing Hoechst. Primary antibody L-plastin (1:500, gift from Yi Feng, University of Edinburgh). Secondary antibody DyLight Alexa 488 (1:250).

### EdU pulse-chase protocol

Larvae of 3 dpf were placed in a 12 wells plate in HEPES buffered (pH = 7.3) E3 containing 0.003% PTU and 0.5 mM EdU for 24 hours. Next, larvae were fixed in 4% PFA at 4°C overnight, dehydrated in 100% MeOH and stored at −20°C for at least 12 hours. Rehydrated to PBS in series and inbated in proteinase K (10µg/ml in PBS) for an hour at room temperature. Followed by 15 minute post fixation in 4% PFA. Larvae were incubated in 1% DMSO in PBS containing 0.4% triton for 20 minutes. Thereafter 50µl Click-iT™ (Invitrogen) reaction cocktail was added for 3 hours at room temperature protected from light. Thereafter samples were subjected to immunolabelling using L-plastin antibody (see section on immunofluorescent labelling).

### Confocal imaging

Intravital imaging in zebrafish brains was largely performed as previously described (66). Briefly, zebrafish larvae were mounted as described for neutral red staining. The imaging dish containing the embedded larva was filled with HEPES-buffered E3 containing 0.016% MS-222. Confocal imaging was performed using a Leica SP5 intravital imaging setup with a 20x/1.0 NA water-dipping lens. Imaging of GFP and L-plastin labelled with Alexa 488 was performed using the 488 nm laser, EdU labelled with Alexa 647 was performed using the 633 laser. Analysis of imaging data was performed using imageJ (Fiji) and LAS AF software (Leica).

### Statistical analysis

For image processing and quantitative analysis SpotNGlia, ImageJ and Prism (Graphpad) were used. Statistical significance was calculated using the one-way ANOVA with Bonferroni correction or Student’s *t*-tests. Standard deviations (SD) are shown as error bars and p<0.05 was considered significant.

---

Supplementary Fig 1. Expression of putative regulators of microglia development in the zebrafish brain.

Bar graphs represent expression values of putative microglia regulators in microglia (green) and other brain cells (blue) observed in the microglia transcriptome (27).

Supplementary Table 1. gRNAs and their mutagenic efficiencies.

## ACKNOWLEDGEMENTS

We thank Mike Broeders for synthesis of high quality Cas9 protein, Esther Hiemstra, Farzaneh Hosseinzadeh, Amber den Ouden, Lucas Verwegen, Ana Carreras Mascaró and Jordy Dekker for their assistance in the reverse genetics screen, and we thank Yi Feng (Edinburgh) for the L-plastin antibody. TVH is supported by an Erasmus University Rotterdam fellowship.

